# Modelling mitosis with multiple phenotypes: relation to Haeckel’s recapitulation law

**DOI:** 10.1101/253203

**Authors:** Yuriy Alexandrov

## Abstract

The article presents a novel stochastic mathematical model of mitosis in heterogeneous (multiple-phenotype), age-dependent cell populations. The developed computational techniques involve flexible use of differentiation tree diagrams. The applicability of the model is discussed in the context of the Haeckelian (biogenetic) paradigm. In particular, the article puts forward the conjecture of generality of Haeckel’s recapitulation law. The conjecture is briefly collated against relevant scientific evidence and elaborated for the specific case of evolving/mutable cell phenotypes as considered by the model. The feasibility, basic regimes and the convenience of the model are tested on examples and experimental data, and the corresponding open source simulation software is described and demonstrated.

## 1. Introduction

Explosive growth of information on molecular components and mechanisms of life constitutes one of the hallmarks of scientific modernity. The main contributors to this process are certainly the biochemistry and biomicroscopy R&D. This study however, only cherry picks some of the major findings of mainstream cell biology to devise the very minimalist computational model of cell proliferation dynamics. The approach is essentially phenomenological and complemented by a corresponding view of evolutionary theory.

The subject of mathematical modelling of cell phenotype dynamics is recurrent in the literature. One of the early but in-depth examples of nonlinear Ordinary Differential Equations (ODE) systems applications to immunology is the work of Perelson et al [1], which actually treats coupled rate equations for the mixture of different cellular and molecular components in blood. More recently, the ODE approach was applied, in particular, to model abnormal regimes in haematopoiesis with multiple phenotypes [2,3], and competition-mediated proliferation in two-phenotype cell culture [4]. It is well-known, however, that the right-hand sides of such ODEs are the subject of meticulous designs bordering with art, as ODEs, being a general method, are not specially suited for cell proliferation modelling. Among stochastic techniques it is worth mentioning the version of the two-phenotype stochastic model of concurrent haematopoiesis [5] extended for the case of several phenotypes [6] and the multi-phenotype model of bone marrow featured by cellular automaton-style considerations [7], the latter being the closest to the presented approach. Another stochastic cellular automaton model by Sundstrom et al [8] studied tumorigenesis with multiple phenotypes by taking into account 2D and 3D spatial effects and limitations, but not cell differentiation. The work by Lu et al [9] analysed the mitosis in two-phenotype systems on the basis of the algorithm by Gillespie [10], also without cell differentiation. In the work by Jones [11], the Bayesian age-dependent branching process modelling was applied to simulate three-phenotype dynamics with differentiation; this method is conceptually quite complex but it does not work with an arbitrary lineage formation tree. An integrated PDE approach was used to devise the two-phenotype model of kidney cell proliferation in a mouse embryo to fit the Optical Projection Tomography (OPT) data [12], and the two-phenotype model of hematopoiesis [13], but these techniques are also hard to generalise for the case of many phenotypes. The same is true for the diverse set of models reviewed in the work on colonic crypt and colorectal cancer modelling [14]. The monograph by Kimmel et al [15] describes several modelling methods applicable to the problem (e.g. the Galton-Watson process); however, these techniques employ a high level of detail and mathematical complexity, likely excessive in many concrete contexts. The work by Stadler et al [16] operates with phenotype-specific parameters similar to those employed in the current study but without focusing on the time dependencies of cell numbers.

Summarising this brief overview of computational methods, one can notice that neither of the existing modelling approaches develops methodology and software specifically for the mitotic multiple-phenotype systems which is both simple, intelligible and convenient for biologists as well as general and flexible enough to cover cross-physiology subjects. This issue is addressed in the modelling chapter, but first the article elaborates on the nature of the mentioned generality.

Namely, the article argues in favour of the abstract Haeckel law for the living (replication-based, evo-devo) systems. The “primordial rock” hypothesis is put forward to emphasize the common evolutionary origin of mitotic cell phenotypes and the general nature of what is vaguely recognized today as “epigenetic determinism” [17], but likely can be attributed to the above mentioned Haeckelian recapitulation.

The next and major part of the study is then devoted to the stochastic, differentiation tree-based mathematical model of heterogeneous mitotic proliferation. The model involves only a handful of essential phenotype parameters, those being cell cycle duration, probabilities of differentiation and exit (which may include apoptosis), and the generation counting threshold. The last property might seem unusual but it actually isn’t. For example, an index identical to the generation counting threshold was introduced in the theoretical model by Jones [11] describing brain cell generation; there, it was explained as “the number of cell cycles [the progenitor] cell must go through before it is competent to produce oligodendrocyte”. In the modelling work on mitosis with cell differentiation by Hasenauer [18], one of the presumptions was that “in most multicellular organisms the mother cell divides symmetrically into two daughter cells which inherit the age of the mother cell”; this “age knowledge” property of cells is also synonymic to a generation counting ability. The biochemical evidence that cells are capable of counting numbers of divisions and utilising this information in decision making, has been recently discovered [19,20].

The article is concluded by examining properties and outcomes of the computational model. The explanatory/predictive capability of the model is confirmed on several test cases, including applications to real-world data on renal development in mice [12] and proliferation of the MDCK cell line in culture [4].

## 2. The Paradigm

As mentioned above, advances in microscopy and in the understanding of genetics in terms of molecular components became defining factors for studying cells at mesoscopic level. In corresponding considerations in the literature, the notions of a cell’s functional “mission” and distinct phenotype are central (e.g. “phenomics”, [21,22]). They reflect not only the existential meaning of physiological purpose, but also common origins and relational directionality - it isn’t possible to get a phenotype from anything else except other (precursor, progenitor, predecessor, parent) phenotypes. It is worth mentioning that the term “phenotype” itself was used in the first place in order to describe physical traits and the morphological/functional sameness of macroscopic animals and plants (albeit later being agglomerations of distinct organs that are in their turn agglomerations of cells). In macroscopic (species)-level biology the term “phenotype” was associated with a whole package of notions dictated by evolutionary principles - mutations, feature heritability, intra- and interspecific competition, “selection pressure”, as well as the well-known phylogenetic evolutionary tree diagrams. Another powerful albeit less frequently mentioned paradigm is Haeckel’s law of embryonic development -” Ontogeny recapitulates Phylogeny”; its basis and meaning have remained a subject of intense theoretical debate for a significant period of time (e.g. [23]). Its modern formulation in zoology is the following: “The definite form of an animal develops on the top of the pre-existing embryonic form of its ancestors” [24]. This formulation implies some sort of directed (in general, one-to-many) relation between ancestral and descendant forms. The general formulation seems even simpler, being almost tautological: “The current biological form develops on the top of the prototype form (of its ancestor)”. This is illustrated schematically in Figure 1. In recent literature, this “develop” keyword becomes more and more associated with the literal, “epigenetic” view of development as a consecutive, ordered activation (expression) of certain groups of genes [17]. The role of genes in recapitulation was also discussed in an earlier work by Ohno [25]. Currently, advances in single-cell transcriptomics facilitate quantitative approaches describing phenotypic diversity in terms of intermediate cell states (ICS) and/or even continua of cellular phenotypes [26]. It is emphasized that every particular change (trajectory) of gene expression patterns is highly constrained while it is being channeled towards state space regions in which all regulatory interactions are “satisfied” (“attractor states”, [27]). One can conclude by generalizing, therefore, that any such trajectory, or in other words, a concerted activation of genes may be (or rather cannot be anything else but) a part of, or an equivalent to “sliding along the branch” of a certain lineage tree of stable ancestral prototypes (“ancestral attractor states”). This seems to agree well with the recent, eloquently articulated statement in the work by Stadler et al [16]: “macro-evolutionary processes happening over million of years may help us to understand microscopic processes on the level of cells”.

**Figure 1.**
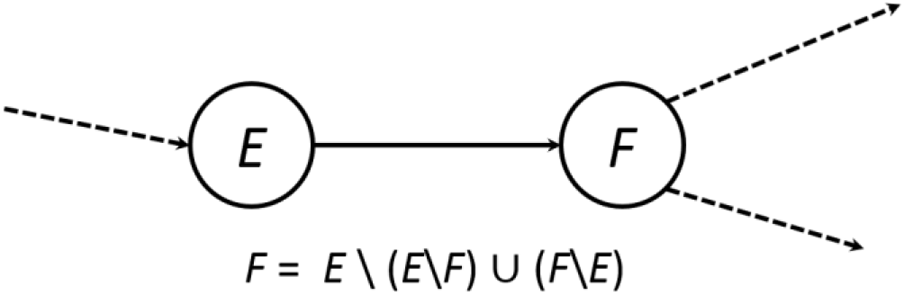
Formal representation of Haeckelian progression from form *E* to form *F*. The “form” (e.g. phenotype) is considered to be a set of well-defined sub-forms. The sets (*E*\*F*) and (*F*\*E*) denote some definite sub-forms developed in *E* but not in *F*, and vice versa. In other words, in order to progress to form *F* having form *E* as a base, one needs to lose some old and acquire some new sub-forms.

Despite many of the evo-devo ideas eventually becoming mainstream in the sub-cellular level literature, the relational tree diagrams for mammalian cell phenotypes had not been studied in great detail until very recently (with the notable exception of haematopoiesis), and the cellular analogue to the Haeckel’s recapitulation law is missing completely. It is interesting to note that in the work by Jones [11], the mathematical formalism developed to model branching processes in molecular evolution (similar to the ones used in the BEAST [28] and MrBayes [29] programs) but also applicable for phylogenetic macroevolution, was adapted to describe progenitor cells’ mitosis and differentiation in the brain, - but in this case the branching tree of cell types was not referred to as “phylogenetic”.

The following statements, albeit hypothetical, are aimed at easing this disparity and hopefully bringing the subject of cellular recapitulation beyond semantics. They also try to elucidate the similarity between cells in terms of the common origin (and common place of origin) of cellular phenotypes.

1. Haeckel’s recapitulation law is the general horizontal principle for all hierarchical levels of life driven by heritability and variation/selection (molecular, organelle, cell, organ, organism).
2. For example, one can consider the well-known haematopoietic lineage tree based on hematopoietic stem cells (HSCs, [30]) as “phylogenetic” in the Haeckelian sense. Haematopoiesis recapitulates the “embryonic” stages of evolution of mitotic pre-multicellular phenotypes in the succession of ancestral liquid organs that eventually became blood. This is the specific (albeit likely the most ancient) case of animal organogenesis.
3. The cellular “evolution” mentioned in (2) once took place in oceanic, shallow-water, calcium-rich geological structures (“primordial rock”, a precursor to coral reefs). The environmental pressure caused by the reefs’ different climatic niches has led to the accumulation of corresponding cell differentiation potentials in their genomes. The internal locus of the bone marrow, known to be the source of blood-forming and of other stem cells in the animal body [31] points at such origin directly. The similarity between the structures of bone and coral is a known fact [32]. It is also known that both calcium dependent bone-forming osteoblasts and bone-resorbing osteoclasts derive straight from bone marrow cells [33]. Yet another hint is calcium signalling, which is ubiquitous in the regulation of such a number of cellular functions, that it may be considered a defining feature of cellular life per se.
4. From this perspective, one can consider the “primordial broth” within that “primordial rock” as the ancestral organ of blood and lymph, and the rock itself as an ancestor of bone tissue.
5. Any cellular process involving proliferation, migration, differentiation and structure formation (e.g. embryo development, tissue regeneration, immune response, inflammation, cancer, angiogenesis, etc.), “plays back” the corresponding pre-recorded elementary steps (patterns) of biotic evolution. The exact triggering mechanism for such “playback” may include different stimuli, such as cell signalling, bacterial infection, injury, etc. In particular, biotic recapitulation is the only way of building a functional supra-cellular level biology (tissue/organ/organism). The lineage tree-driven separation of proliferating phenotypes in time is in many cases (rather most, for solid tissues) followed by a concomitant separation in space. The well-confirmed HSC “niche” concept [34] in that matter does not differ from any other organ-specific stem cell pool or niche.
6. The phenomenon known as cancer is a specific case of organogenesis, implementing pre-recorded phenotype progression routes that are, arguably, either archaic, or concomitant (parasitic) to mainstream lineage formation. In viable organisms, these biogenetic routes are either blocked or have negligible probability, but they may be initiated e.g. by mutation.
7. During development via Haeckelian recapitulation, cell populations are controlled by generation counting. Namely, the precursor phenotype cells count the number of divisions during mitosis and, after reaching some threshold number, progress to the (one of the) next allowed phenotype(s). It is reasonable to assume that such generation-counting control is behind organ scaling and allometry phenomena [35]. The evidence of as well as the prospective mechanisms and biochemical basis for cell division counting have recently been discovered [19,20]. In particular, in algal cell cultures, this counting mechanism is associated with a specific protein CDKG1 [19]; the amount of this protein in the cell is reduced after each division.

## 3. Computational model

The computational recipes listed below do not essentially rely upon any of the potentially questionable aspects of the “paradigm” presented above, except for possibly the generation counting principle. However, the latter can be considered from a purely pragmatic viewpoint as well, by simply treating generation counting thresholds as adjustment parameters of the model. On the other hand, taking “The Paradigm” into account can be beneficial for further progress in understanding the evolutionary “logic” behind every concrete case of recapitulation-driven proliferation.

The main components and principles of the computational model are:

- Distinct cell phenotypes capable of differentiating from one to another.
- Stochasticity. This is essentially stochastic technique.
- Cell cycle. In the code, the corresponding parameters are the cell cycle duration *T*_*c*_ and its standard deviation *ΔT*_*c*_ ; they are different for different phenotypes.
- Differentiation lineage trees, implemented as directional graphs in the code, with every node representing a phenotype, and every edge representing the probability of differentiation from the current to the next allowed phenotype.
- Direct control of cell numbers by the exit probability.
- Generation-iterating method (“God’s algorithm”, see below).
- Generation-counting thresholds.

Among these principles, only the last two are novel, and as such require further specification. The generation-iterating method is recruited as a shortcut to circumvent the difficulty of iterating the time-dependent state variables in stochastic modelling. If one chooses some fixed *Δt* as a time increment driving physiological dynamics, all results become dependent on the value of *Δt*. Using rate-dependent methods such as the Gillespie algorithm (Gillespie 1984) to overcome this adds significant mathematical and computational complexities. In the present study, the problem is avoided by using the observation, that in some cases, having a set of (g)-generation cells is fully sufficient for deducing the set of next (g+1) generation cells. In the simple implementation presented in this article, for every newly created (g+1)-generation cell, the following parameters are immediately defined:

1. If it exits the pool or not, instead of continuing (according to exit probability *α*)
2. The phenotype - either the parent’s, or the next (depending on the generation count threshold)
3. Generation count - either the parent’s+1, or 1 (ditto)
4. Birth time - equals the end of the life time of the parent cell
5. End of life time - (birth time + randomized *T*_*c*_, depending on the phenotype)

During simulations, properties 1, 2, 3 and 5 are stochastically chosen for every new cell, whereas properties 2, 3, 4 and 5 are actually kept in computer memory. The term “God’s algorithm” reflects the usage of property 5, which implies “knowing the future” of a cell’s fate at the time of its birth. Herewith, the cell lifetime values are randomized by sampling from a Gaussian distribution centred at *T*_*c*_ with standard deviation *ΔT*_*c*_. In order to estimate the number of cells, *N*(*t*), as a function of time *t* from such simulations, one needs to count those cells for which time *t* falls within their lifespan (Figure 2). The drawback of the algorithm is that in order to do this, one needs to keep a complete record of the history of all cells in memory. Fortunately, this is not very problematic, since as was shown for the simplest case above, every cell is represented by only 4 numbers (type, generation and the birth and end of life times). Its advantage is that it allows replacing iteration by time by the iteration of generations.

**Figure 2.**
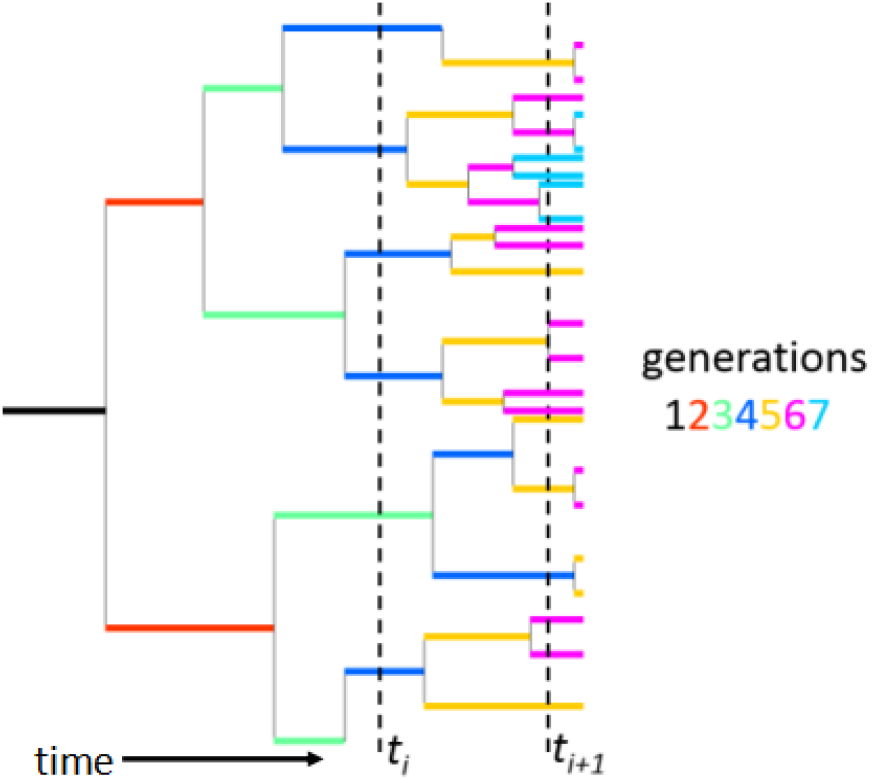
“God’s algorithm” in action. Horizontal segments represent cells of different generations, correspondingly colour-coded. When computing the time dependencies of cell numbers, one needs to count the number of segment intersections (‘hits’) made by the sampling *t*-lines (dashed black vertical lines)

The influence of exit probability *α* on cell proliferation dynamics can be illustrated by the following considerations. Imagine *N*_*0*_ cells of the same type, starting their divisions at time *t* = *t*_*0*_ with average cell cycle duration *T*_*c*_. If *α* has the meaning of the fraction of cells leaving the pool after each division, then the expected number of cells as a function of time, *N*(*t*), can be written as

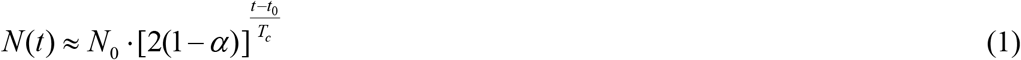

Formulae similar to (1) have been published in the literature (e.g. [12], [36]). It is worth noting that in expression (1), the two essential parameters *T*_*c*_ and *α* are competing in the proliferation rate, since 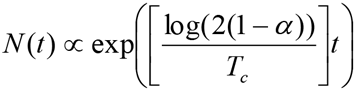. It is therefore important to have reasonable estimates of the cell cycle durations *T*_*c*_ when specifying the model. These can in many cases be provided by microscopy (e.g. [4],[12]).

The last feature of the framework is the utilisation of generation counting thresholds. This is also straightforward. Certainly, having the proper number of intermediate proliferating phenotypes is critical, as their influence is amplified exponentially via mitosis. This is illustrated in Figure 3, where the corresponding cell numbers at threshold level are shown to be 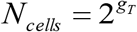, where *g*_*T*_ is the generation counting threshold (number of divisions the cells of this phenotype, on average, perform). In the implementation, not only the generation threshold *g*_*T*_ is taken into account, but also its dispersion *Δg*_*T*_, which represents the degree of stochasticity in the model’s decision-making. Both parameters are fed into a standard sigmoid expression to get the corresponding probability of progression to the next allowed phenotype. Thus, for a *g*-th generation cell, this differentiation probability *P*_*diff*_(*g*) reads as

**Figure 3.**
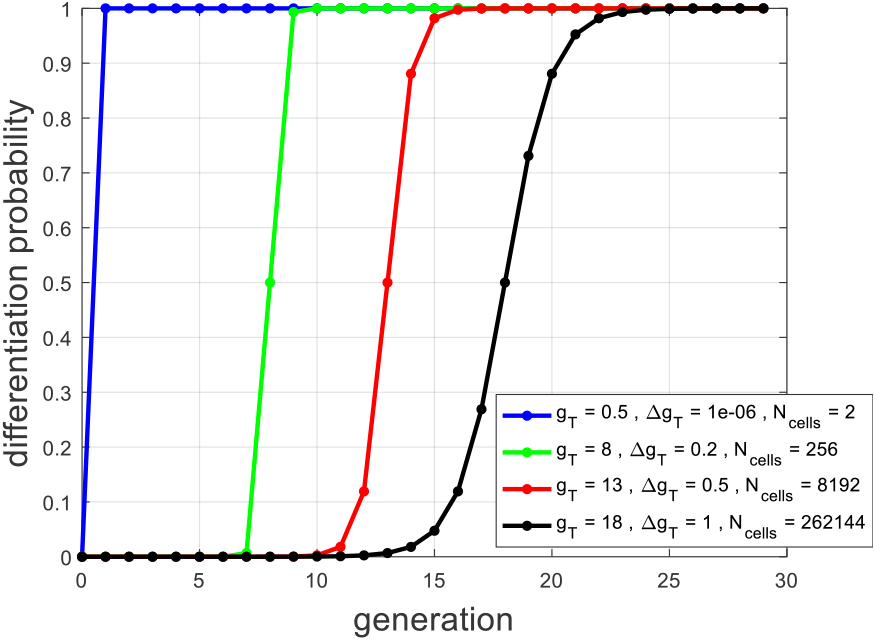
Sample phenotype progression probability dependencies as a function of local generation count, ranging from threshold-like transitions (blue) to smooth ones (black)

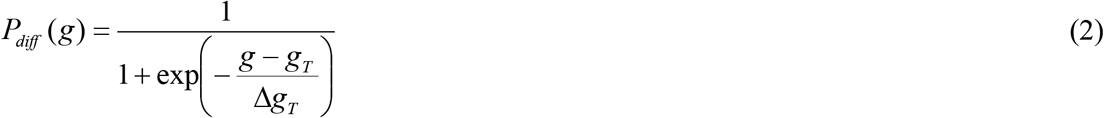

In simulations, the probability *P*_*diff*_(*g*) is used with a uniform, unit-interval distributed random variable to get a randomized “yes/no differentiation” response at the moment of every cell’s birth.

Altogether, every phenotype in the model is characterized by 5 parameters (*T*_*c*_, *ΔT*_*c*_, *g*_*T*_, *Δg*_*T*_, and α), plus the out-degree (the number of outcoming edges) of the node in the directed lineage tree. Every edge in a tree is characterized by the weight representing phenotype-to-phenotype transition probability. For each node therefore, the weights of the outcoming edges sum up to 1. In the course of simulations, if the stochastic choice (made at the moment of cell birth with probability *P*_*diff*_(*g*) given by the equation (2)) favours cell differentiation, the cell differentiates into one of the types defined by the corresponding lineage tree edges outcoming from the parent node. Herewith, the next specific phenotype of a cell is defined by a roulette wheel method involving the weights of a preceding phenotype’s outcoming edges.

It is quite obvious, that any change in the “flow of cell numbers” along the branches of the differentiation tree diagram will eventually lead to the formation of cell populations of differing size (as represented by final phenotypes), and therefore will cause effects such as organ scaling and allometry in embryo development. In the absence of exit, these “flows” of cell numbers from a given node are controlled by the weights of the outcoming edges and by the parameters of generation count thresholding in (2) responsible for the total number of cells of that node’s phenotype.

Finalizing this section, it is relevant to note that the model allows reproducing biologically meaningful population dynamics on the basis of an arbitrary lineage tree without resorting to complex mathematics like the Bayesian inference. On the other hand, the model is neither limited by the specifically chosen proliferation method (iterating generations), nor by the absence of the dependence of cell phenotype parameters on other factors participating in adaptive control such as cytokines and growth factors. The numerous and sophisticated models of cell sensing, interaction and competition have already been known for a relatively long time (e.g. [1]). Such improvements and extensions of the technique are possible and can be investigated in the future.

## 4. Model implementation

The model was coded in Matlab (Mathworks, Inc.). To represent and use differentiation lineage trees, ***digraph*** and related graph functions were used. A simple, intuitive GUI is provided. The computational performance of the model is satisfactory for simple cases involving about 10^4^ cells. However, when the total number of cells exceeds 10^6^, the algorithm slows down noticeably. No attempt has been made to improve performance via parallelising, although it seems possible. The software is available at https://github.com/yalexand/CellPopSim.

## 5. Model properties, results and conclusions

Typical dynamic regimes can be examined with simple linear succession graphs, containing just 2 to 4 consecutive nodes. In doing so, the overall validity of the model is assessed, which is shown is the following. As to the basic properties, the model demonstrated the expected behaviour. Simulations usually start from a single initial proliferating cell of an initial phenotype. In the demonstrative examples drawn below, parameters *T*_*c*_ and *ΔT*_*c*_ are set for all phenotypes within a range of 15-25 ± 2h, which is typical for eukaryotes’ mitosis.

Figure 4 presents an example of a linear, three-phenotype succession showing two types of exponential dynamics for the final phenotype *E*: “disappearance” at exit probability *α*>1/2 (Figure 4, left) and unrestricted growth at *α<*1/2 (right).

**Figure 4.**
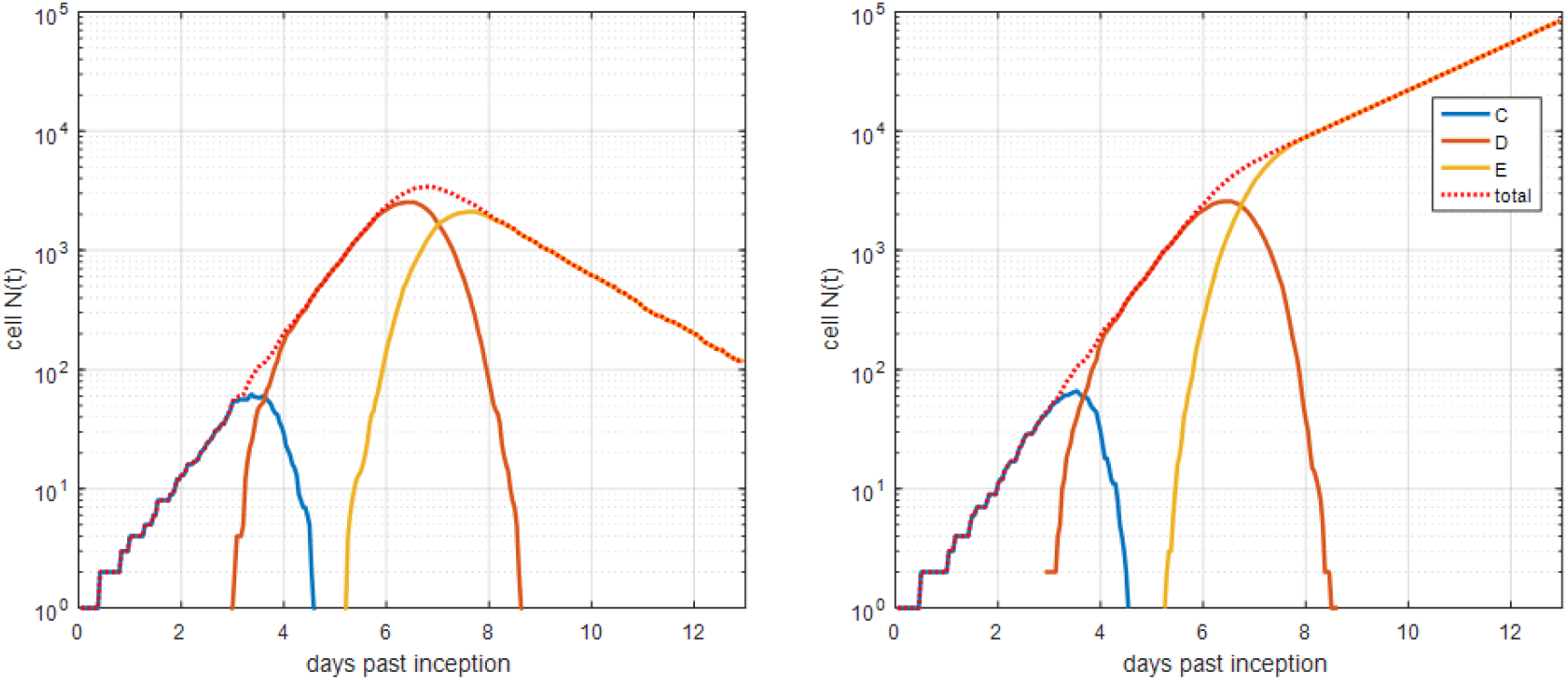
Sample linear three-phenotype succession *C*→*D*→*E* showing two types of exponential dynamics for final phenotype *E*: “disappearance” at exit probability *α=0*.*65* (left), and unrestricted growth at *α=0*.*35* (right)

The Figure 5 compares two ways of reaching equilibrium for the final phenotype. In the first, mitosis continues indefinitely with *α*=1/2 but also with *g*_*T*_=∞ (exit-mediated equilibrium). The second is establishing genuine quiescence at *α*=0, *T*_*c*_=∞, in which case *g*_*T*_ becomes irrelevant (Figure 5 left and right, correspondingly). The higher level of the plateau on the right plot is due to the *α*=0 setting, which preserves all created *E*-cells.

**Figure 5.**
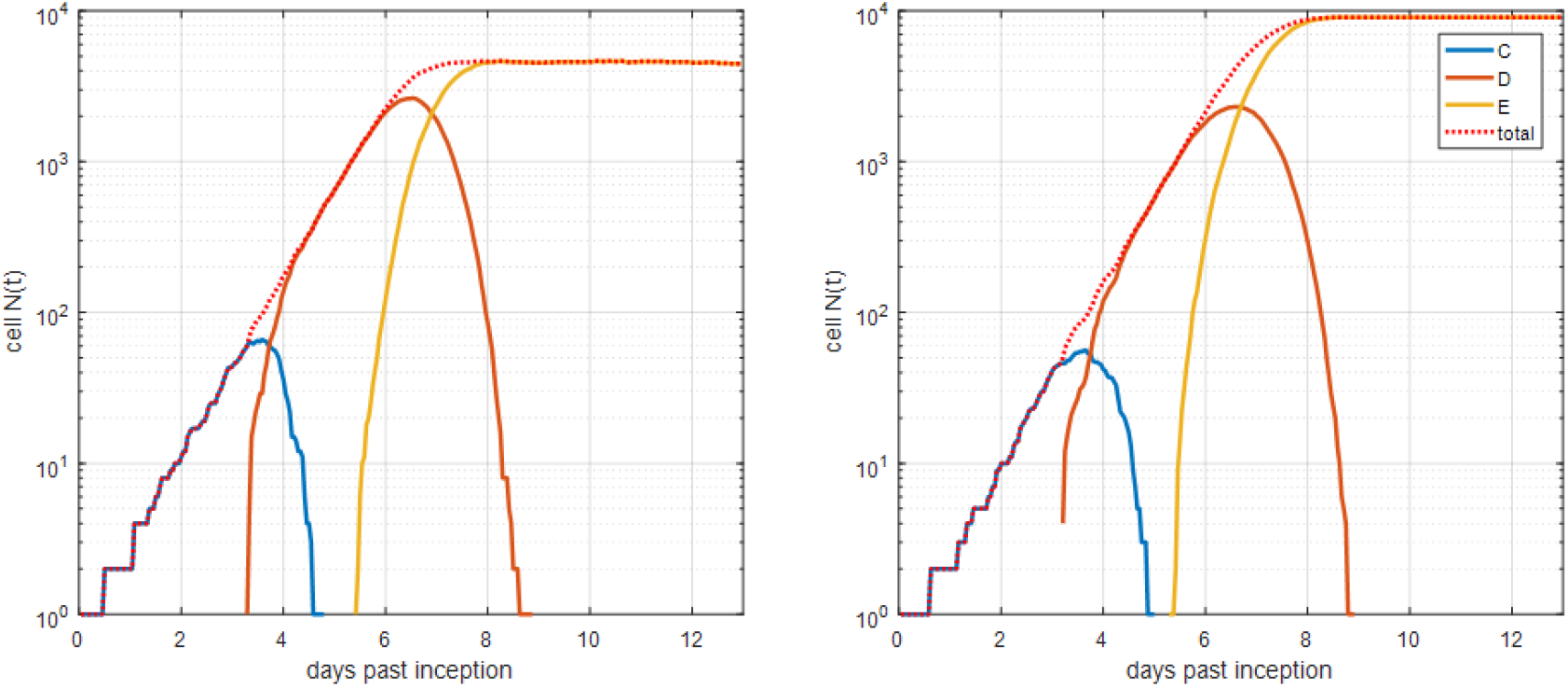
Sample linear three-phenotype succession *C*→*D*→*E* showing the two types of “stable state” for final phenotype E: “dynamic” at exit probability *α=0*.*5* (left), and genuine quiescence at *α=0, T*_*c*_*=∞* (right)

The next example presents the four-phenotype linear succession which keeps overall exponential growth (Figure 6, left), and the same system with a higher (*g*_*T*_=10 vs. 3 for the plot on the right) generation count threshold and an *α*=1/2 setting for the phenotype *D* (vs. *α*=0 for the plot on the right). The *α*=1/2 setting removes exactly half of the newly born phenotype *D* cells during proliferation. Altogether these settings for phenotype *D* allow delaying the exponential growth of phenotype *F* for about 4 days (Figure 6, right). Therefore, such an intermediate phenotype with *α*=1/2 and higher *g*_*T*_ can serve as a “carrier to the future” for subsequent phenotypes, providing the needed flexibility, e.g. in the case of delayed organogenesis.

**Figure 6.**
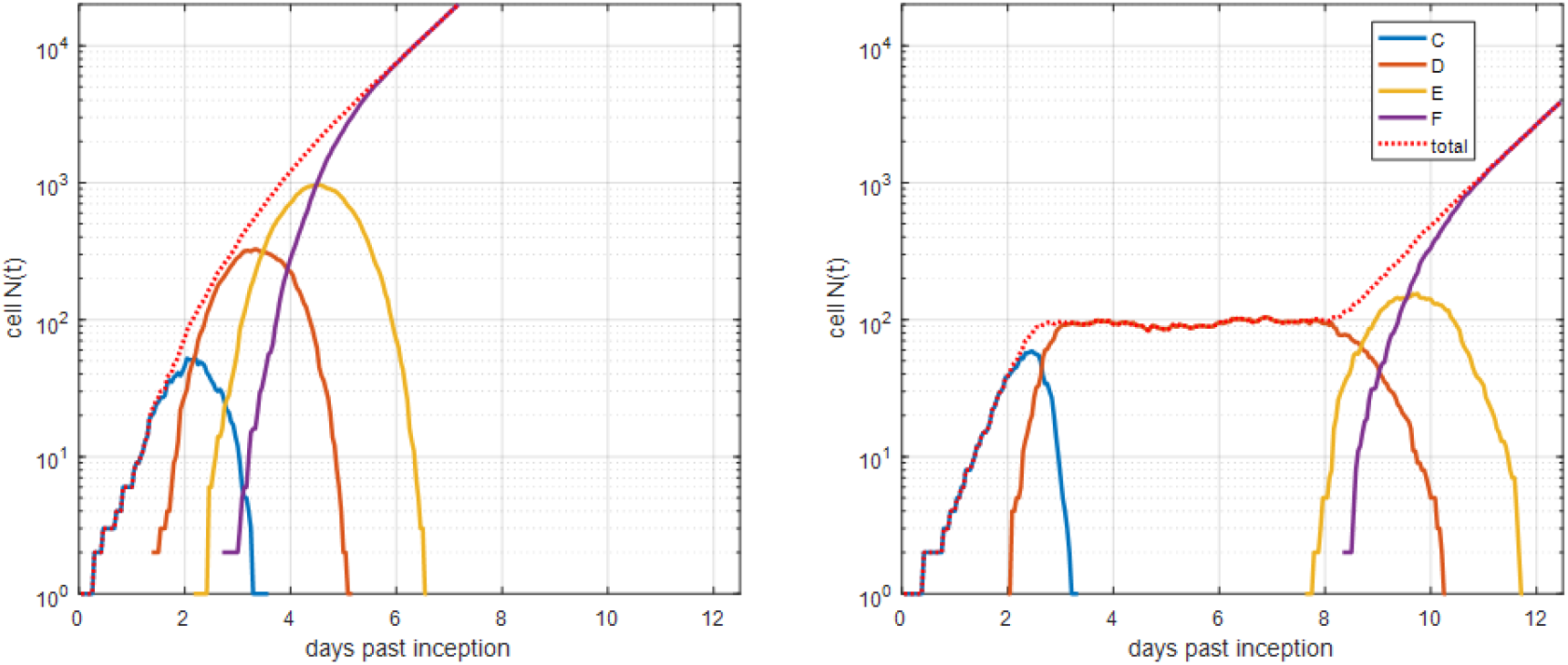
Sample linear four-phenotype succession *C*→*D*→*E*→*F* showing unhindered exponential growth (left), and delayed “*α*=1/2 carrier”-mediated growth (right)

The simple, six-phenotype forked proliferation “outburst” is shown in Figure 7. On the graph to the left, the terminating nodes (final phenotypes) *E, G* and *A* are all “dumped” to decay by setting *α*=0.8. On the graph to the right, the differentiation probabilities for nodes *G* and *A* (edges *F*→*G* and *F*→*A*) are swapped, resulting in their curves changing places, whereas the *E* node was made the quiescent “single survivor” by the setting *T*_*c*_ = ∞. The bar plots display the relative number of cells of different types in the studied time window, which makes sense for the case of “outburst”.

**Figure 7.**
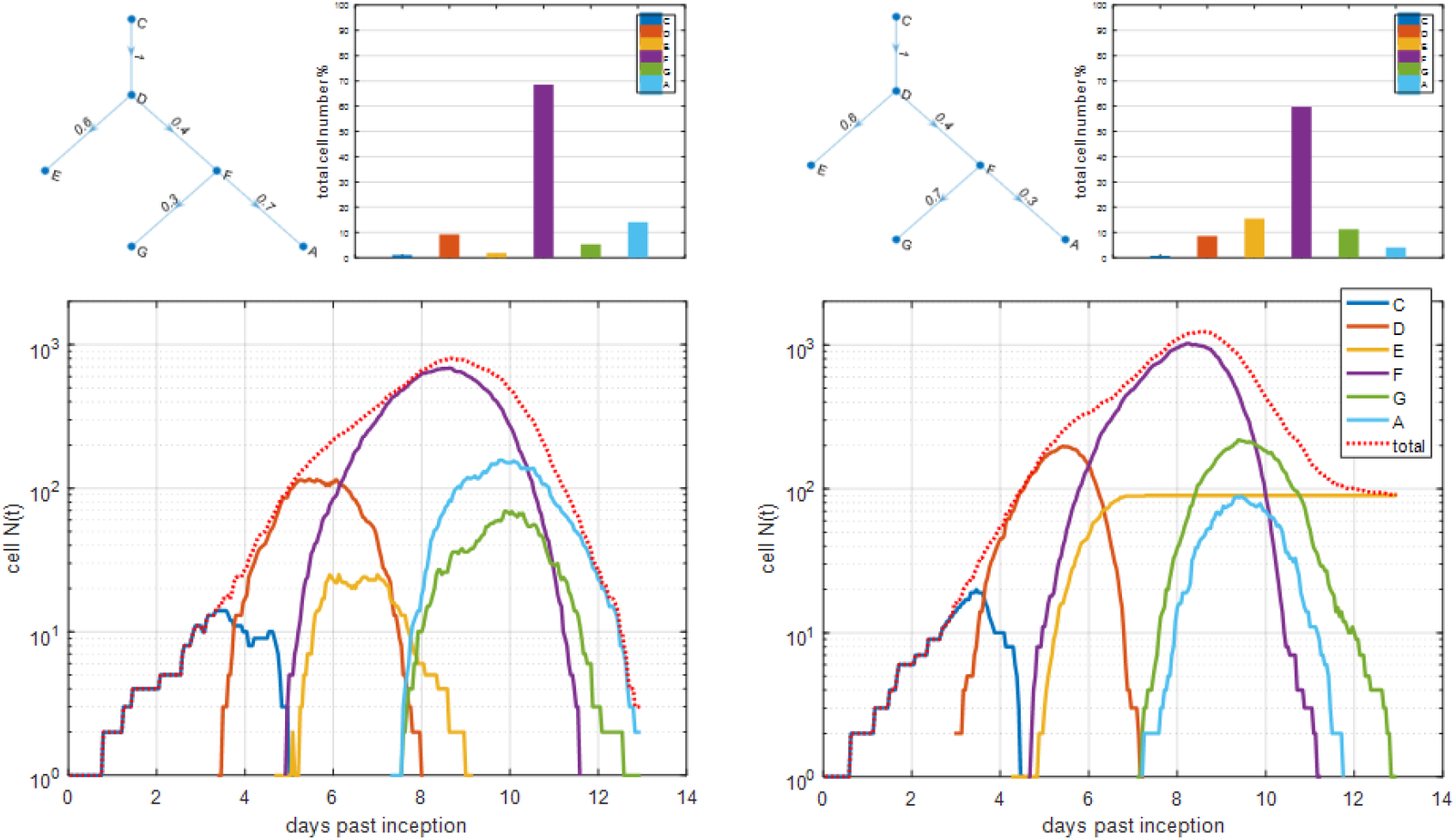
Sample forked five-phenotype proliferation “outbursts”. The terminating nodes *E, G* and *A* are all subjected to exit probability *α*=0.8, which provides fast decay (left). On the graph to the right, the differentiation probabilities for nodes *G* and *A* (edges *F*→*G* and *F*→*A*) are swapped, resulting in their curves changing places, whereas the *E* node was made the quiescent “single survivor” at *T*_*c*_*=∞*

Another example presents the phenomenon of undulations initiated in the seven-phenotype “outburst”-type branching system (Figure 8, left) when one of the branching nodes is assigned a self-renewing property whereas all other parameters stay the same (Figure 8, right). The undulations develop due to the interplay between cell cycle duration *T*_*c*_ and the generation-counting threshold *g*_*T*_, which demands the periodic reset of the generation counter to *g*=1 for the self-renewing phenotype. Certainly, if *g*_*T*_<1 and *Δg*_*T*_<<*g*_*T*_, the only two generations allowed for this phenotype are either *g*=1 or *g*=2; this is the way to effectively “undo” generation counting. It is hard to say if a self-renewing node with *g*_*T*_ >1 (as in Figure 8, right) can represent any biological reality. Nevertheless, as this explanation shows, such hypothetical effects of “self-renewal undulations” at *g*_*T*_ >1 are intrinsic to the model. On the other hand, the involvement of many unrelated proliferating sources will cause undulation smearing, in which case undulations will not be noticeable in the experimental data.

**Figure 8.**
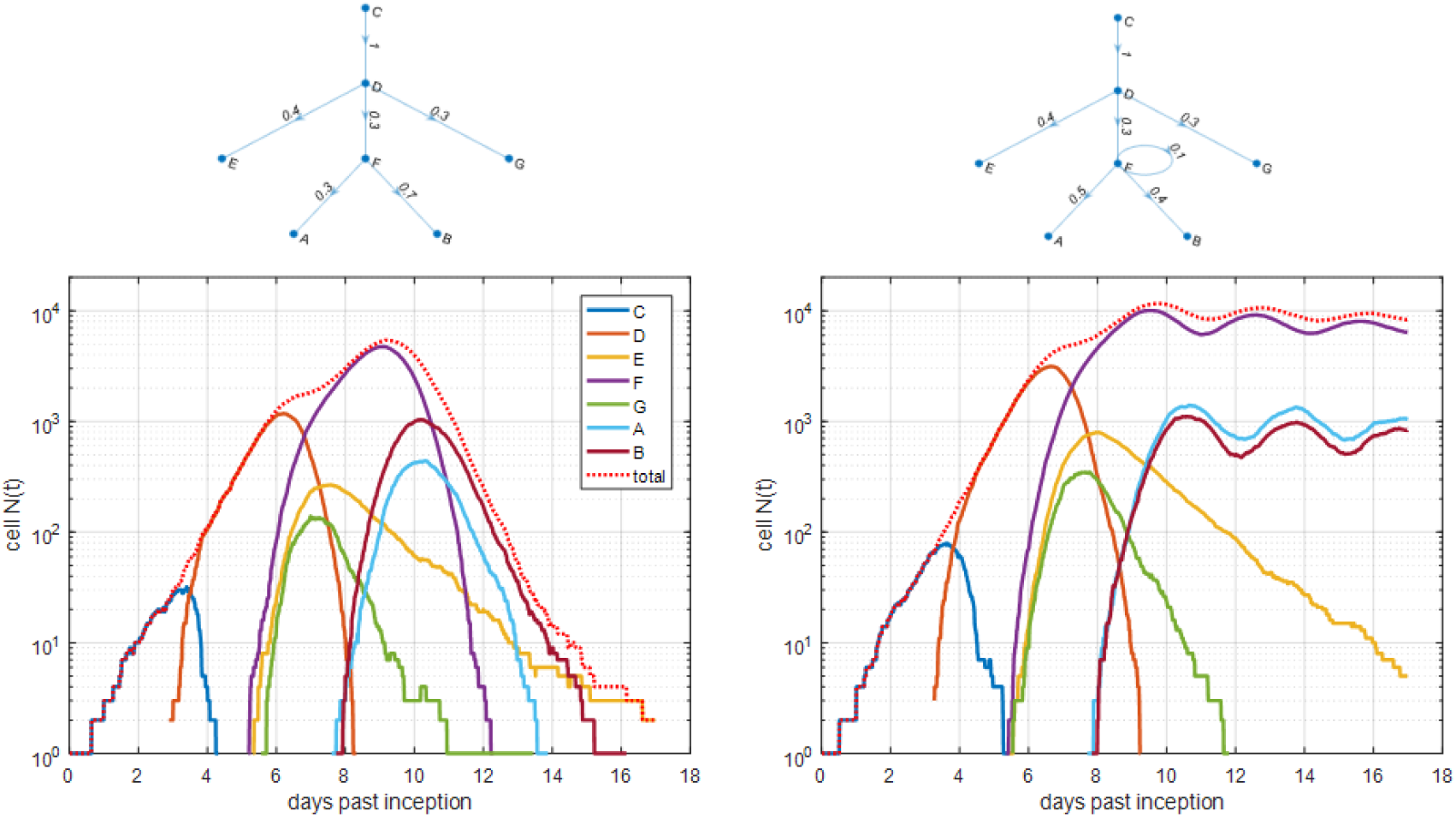
Sample forked seven-phenotype proliferation progressing from “outbursts” (left) to undulations (right) after the addition of a self-renewal property to node *F*. Similarly to the example shown in Figure 6, terminating nodes *E, G, A* and *B* are all subjected to exit probability *α*=0.8

Another example of undulations, this time caused by introducing the de-differentiation edge, is shown in Figure 9. In this case, the generation counting was “undone” for both involved nodes as described above, by setting *g*_*T*_<1 and *Δg*_*T*_<<*g*_*T*_, so that the period of such undulations is defined only by the cell cycle duration values, *T*_*c*_, of these nodes. Undulations also develop when terminating nodes are involved (data not shown). It is relevant to note that the undulations of cell numbers have been reported in haematopoiesis data and studied in corresponding models [2,3].

**Figure 9.**
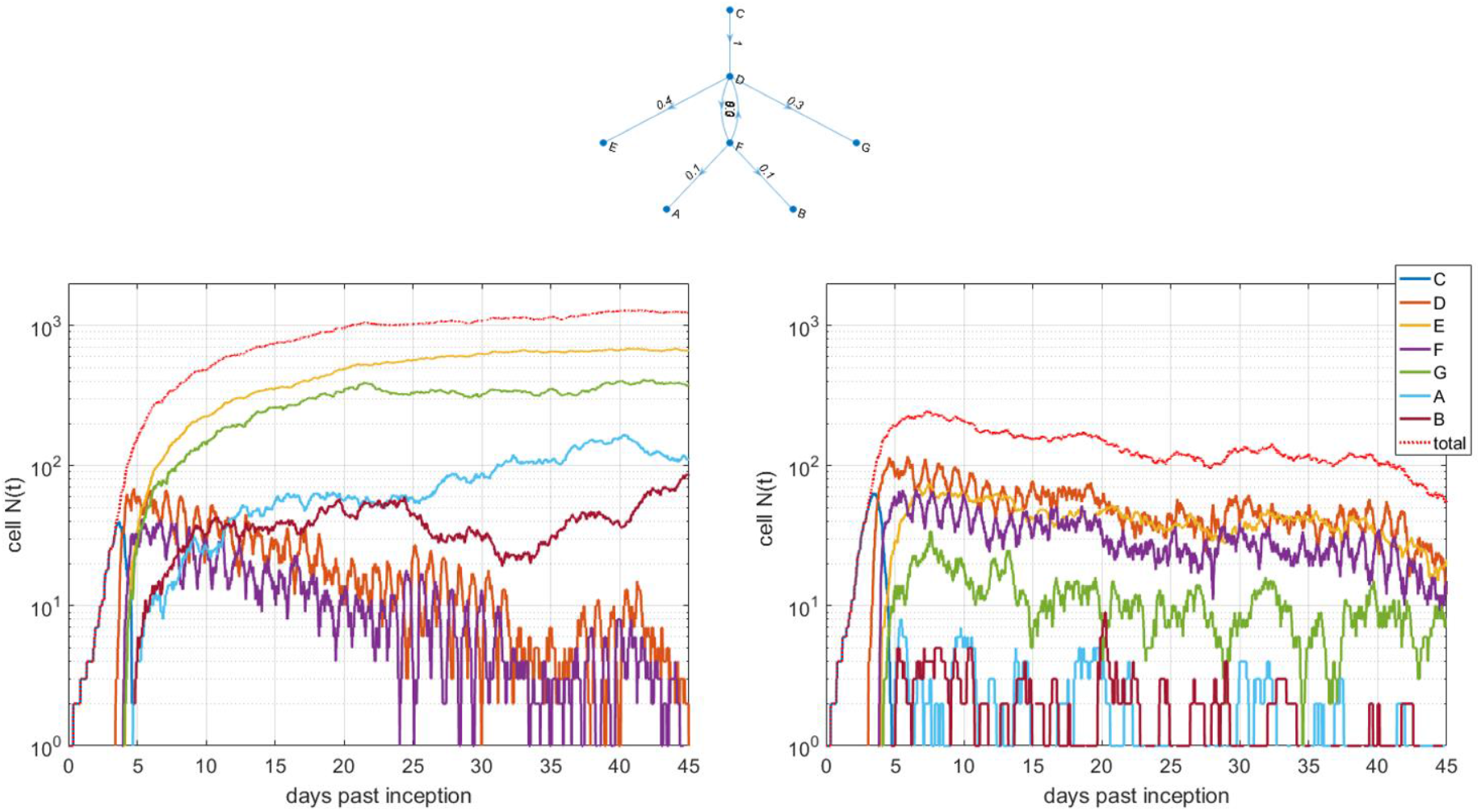
Sample forked seven-phenotype proliferation progressing from a “stable state” where the terminating nodes *E, G, A* and *B* are all subjected to exit probability *α*=0.5 (left) to decaying “outbursts” at *α*=0.8 (right). In both cases, the undulations in *F* and *D* cell numbers are caused by de-differentiation introduced by edge *F*→*D*

The next example (Figure 10) represents the application of the model to the experimental OPT data reported in the article on mouse kidney development by Lefevre et al [12]. This study reports a very steep near-exponential growth of the mass of the precursor cap mesenchyme (CM) cells at about 2 weeks past conception, before differentiation to the final renal ventricular (RV) cell type. In the article, the volume of the CM cells was reported in voxels, with 1 voxel = 4.17µm^3^. Assuming a cell radius of a≈6µm, the estimate for cell volume is (4/3πa^3^)/4.17≈217[vox], and therefore Data[vox]/217 should be close to CM cell numbers. The first “ini” phenotype is used as an adjustment. The article reports two CM sub-phenotypes at around 2dpc with *T*_*c*_ =19.6h and *T*_*c*_ =15h and comparable amounts, which was implemented in the model. The “slow CM” is then used as a “carrier” precursor to the point where the fast growth of CM cells starts (“final CM”). The resulting adjustment for this phenotype was done for the “no apoptosis” case, *α*=0, as suggested in the article, and *T*_*c*_ =18h. The RV phenotype’s settings included *α*=0 and *T*_*c*_ =∞. As shown in Figure 10, the model allows presenting the appearance of the final RV phenotype at a very early stage of renal development and in a quantitative manner. The more rigorous application of the technique can involve parameter estimation and uncertainty analysis, as well as larger experimental datasets (as compared with the number of model parameters); however, this is beyond the scope of the current study.

**Figure 10.**
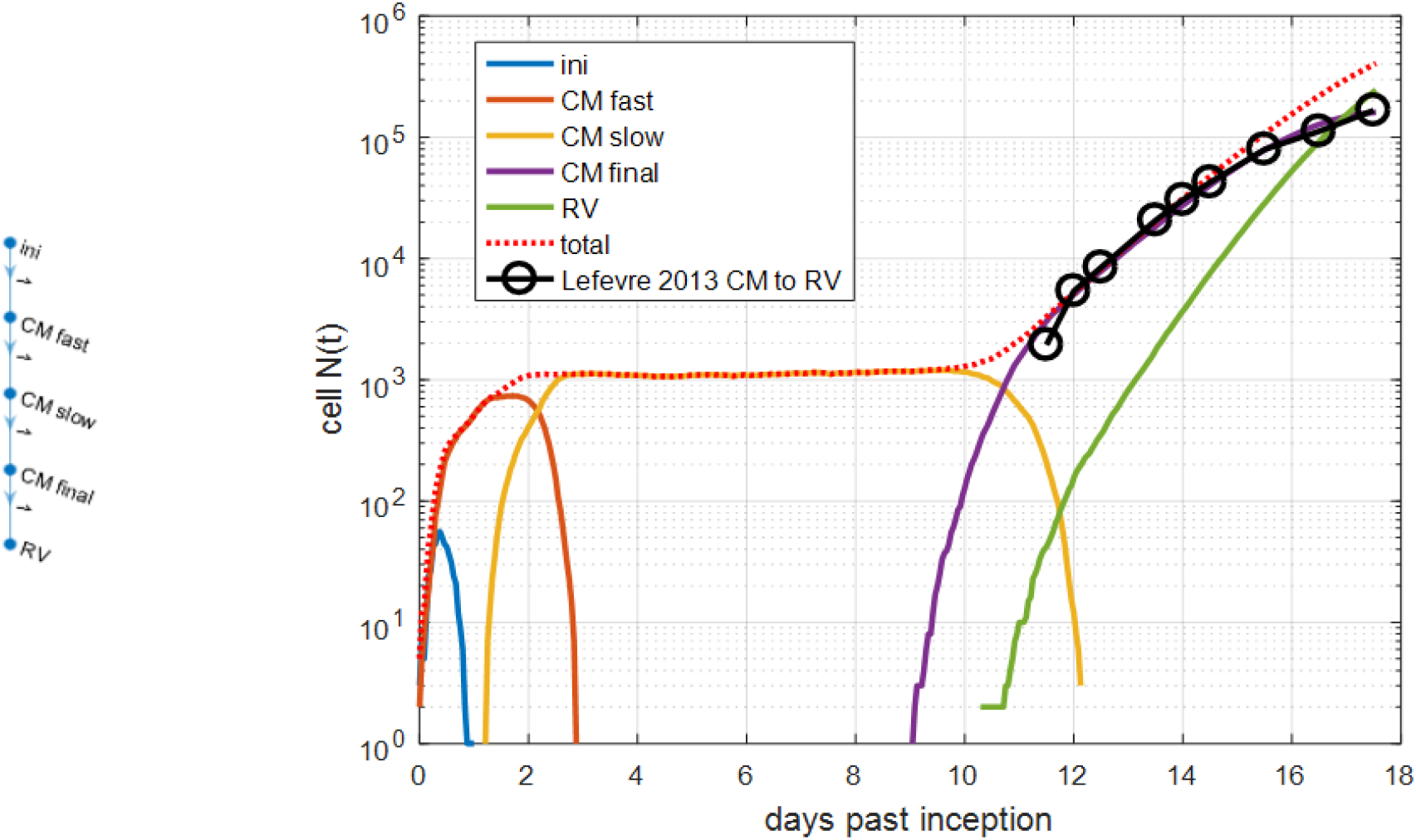
Application of the model to renal organogenesis by fitting OPT microscopy data published by Lefevre et al [12]. The data are used with the authors’ permission. Full details in Section 5

The last example considered in the study is the volume-limited proliferation of the immortalized MDCK cell line in a Petri dish, reported in a study of homeostasis with cell competition [4]. The competing phenotype was represented by the same MDCK cells, but lacking the polarity protein scribble. The present study, however, focuses on the basic (wild type) MDCK^wt^ cells only (the green curve in Figure 2f of the article). The MDCK^wt^ cell numbers, as a function of time (Figures 11 and 12) and cell cycle duration *T*_*c*_ =18h were measured with very high precision. As shown in the Figure 11, first, the approximately exponential middle part of the dependency of cell number on time is chosen in order to fit it to expression (1); fitting was performed with the cell cycle duration fixed at *T*_*c*_ =18h as suggested in [4]. Fitting was carried out by a multidimensional nonlinear minimization (Nelder-Mead, [37]). This allowed estimating the apoptosis probability, *α*≈0.056, and the starting cell number, *N*_*0*_. Next, these parameters were used to define the “seed” and the middle “mitotic” phenotype in the simple chain proliferation model with three “phenotypes” termed “seed” (proliferating, with *α*=0 and *g*_*T*_<1), “mitotic” (proliferating with *T*_*c*_ =18h and *α*=0.056) and “inhibited”, respectively (Figure 12). It turns out, that in order to find a good adjustment for the whole dataset, it is enough to set the apoptosis rate of the “inhibited” phenotype to the familiar value of *α*=½. In the experiment and the article’s discussion, these three “phenotypes” were certainly not distinguishable from one another. However, this example, as well as the previous one, demonstrate that such decomposition of cells into groups according to their “mission” can be applied in a variety of cases to describe hypothetical population dynamics. In particular, it allowed interpreting volume-controlled proliferation with the help of a new, “inhibited” phenotype.

**Figure 11.**
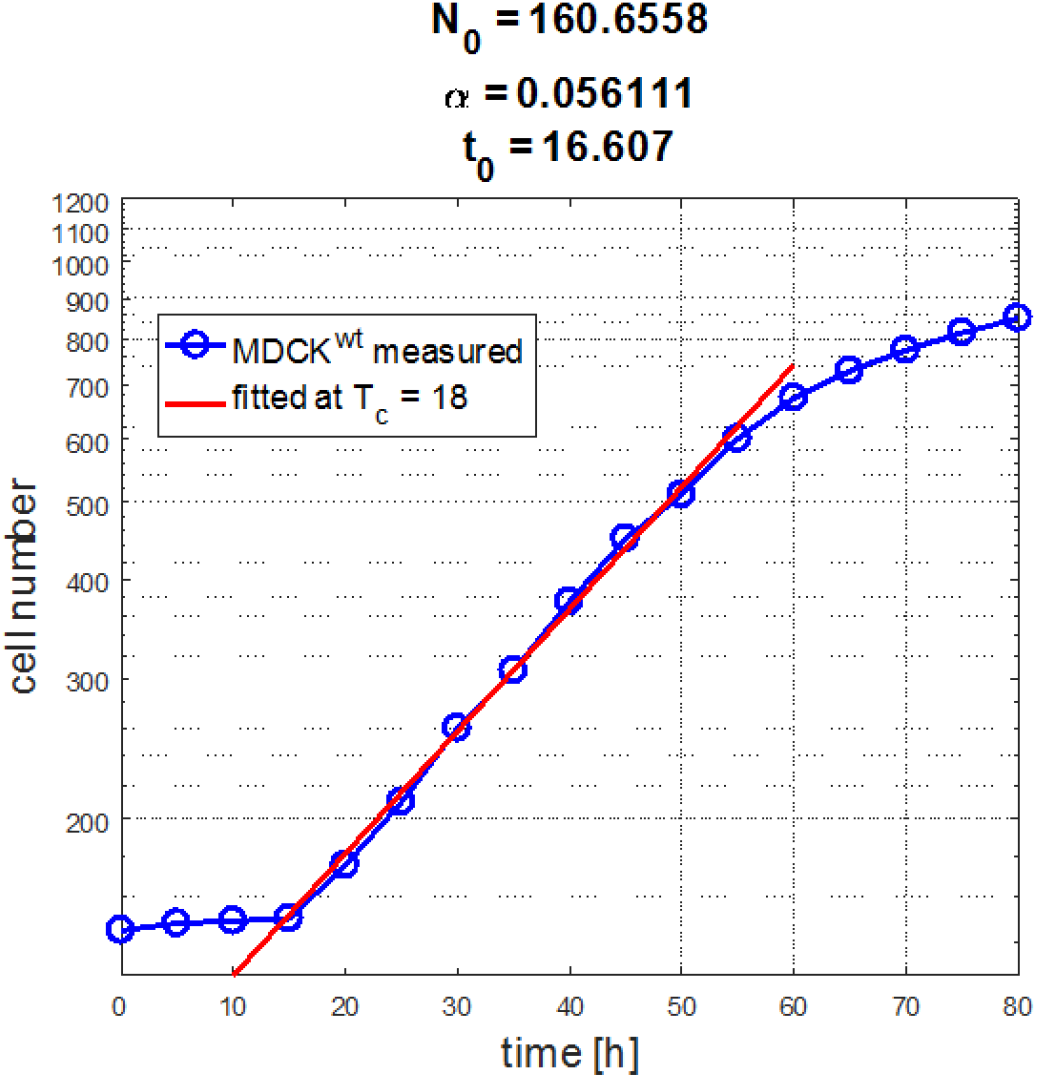
Fitting of the time dependence of the number of proliferating MDCK cells published in Bove et al [4] to expression (1), performed using the Matlab *fminsearch* function. Note the log Y scale. The data in Fig 11, 12 are used with the authors’ permission

**Figure 12.**
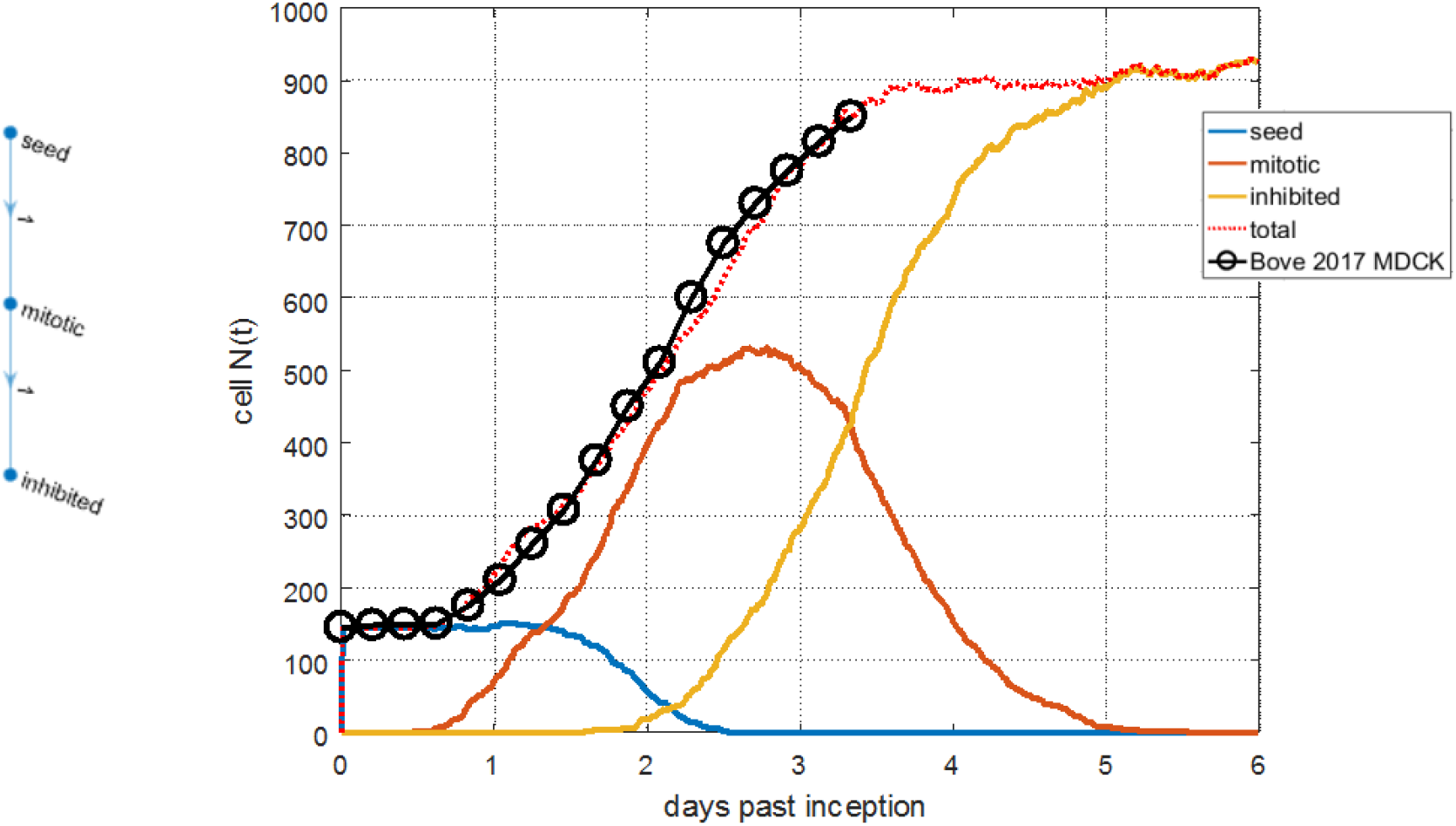
Adjustment of the time dependence of the number of proliferating MDCK cells published in Bove et al [4] using a three-stage proliferation model

Coming back to the question of parameter estimation, one should note that from this viewpoint, the two cases of model application to real-world samples considered above (CM/RV and MDCK) only present a single stochastic iteration. For a given case, strictly speaking, that is not enough to infer the role and/or argue in favour of the sufficiency of the generation counting mechanism. The latter, however, when being implemented as a control technique, definitely provides for the “rigidity” of the developmental program, a feature associated with the concept of “attractor states” [27] and certainly with biological reality per se. Indeed, the very “rigidity” of embryonic development has defied mechanistic explanations since Haeckel’s times, but it arises naturally in the proposed method. The thorough investigation of model parameter space remains a challenge for future research, as these iterations are complex and computationally demanding.

In conclusion, it is worth emphasizing again that the motivation and intrigue behind the developed computational model of cell proliferation was the conjecture of the generality of Haeckel’s recapitulation law as propounded in “The Paradigm” section. According to this conjecture, Haeckel’s law is one of the overarching principles of biological causality. The presented model shows how it can contribute to the understanding of mitotic proliferation with multiple differentiating phenotypes. Although the exact molecular mechanisms of genetics-driven recapitulation have yet to be discovered in full, it is believed that with the presented approach, coupled with the support of viable mathematical models, mechanistic “recapitulation physiology” can be introduced and further utilised to the benefit of biological R&D as a whole.

## 6. Acknowledgments

The author thanks Mr. Oleg Alexandrov for revising the English of the manuscript. Author gratefully acknowledges partial funding from the UK Medical Research Council (MRC, MR/K015834/1), the Biotechnology and Biological Sciences Research Council (BBSRC, BB/M006786/1).

